# Occurrence of non-apical mitoses at the primitive streak, induced by relaxation of actomyosin and acceleration of the cell cycle, contributes to cell delamination during mouse gastrulation

**DOI:** 10.1101/2024.01.24.577096

**Authors:** Evangéline Despin-Guitard, Steffen Plunder, Navrita Mathiah, Eric Theveneau, Isabelle Migeotte

## Abstract

During the epithelial-mesenchymal transition driving mouse embryo gastrulation, cells at the primitive streak divide more frequently that in the rest of the epiblast, and half of those divisions happen away from the apical pole. These observations suggests that non-apical mitoses might play a role in cell delamination and/or mesoderm specification. We aimed to uncover and challenge the molecular determinants of mitosis position in the different regions of the epiblast through a combination of computational modeling and pharmacological treatments of embryos.

Blocking basement membrane degradation at the streak had no impact on the asymmetry in mitosis frequency and position. By contrast disturbance of actomyosin cytoskeleton or cell cycle dynamics elicited ectopic non-apical mitosis and showed that the streak region is characterized by local relaxation of the actomyosin cytoskeleton and less stringent regulation of cell division. These factors are essential for normal dynamics at the streak but are not sufficient to promote acquisition of mesoderm identity or ectopic cell delamination in the epiblast. Exit from the epithelium requires additional events, such as detachment from the basement membrane.

Altogether, our data indicate that cell delamination at the streak is a morphogenetic process which results from a cooperation between EMT events and the local occurrence of non-apical mitoses driven by specific cell cycle and contractility parameters.

## INTRODUCTION

Epithelial-mesenchymal transition (EMT) is a reversible process by which a static epithelial cell becomes motile through the acquisition of a mesenchymal phenotype. It involves a loss or change in intercellular junctions, reorganization of the cytoskeleton, a switch from apical-basal to front-rear polarity, and remodeling of the extracellular matrix^1,2^. EMT is a fluid non-linear spectrum in which cells can adopt intermediate phenotypes with various levels of epithelial and mesenchymal features^2^.

Gastrulation, an evolutionary conserved developmental event through which multiple germ layers arise from a single epithelium, occurs through EMT-mediated cell delamination. In the mouse embryo, gastrulation takes place in the primitive streak (PS), a structure specified in the posterior region of the epiblast at embryonic day (E) 6 ^3^.

One of the earliest steps of gastrulation EMT is the degradation of the basement membrane^4,5^. Cells maintain tight and adherens junctions until they exit the epiblast^3^, then their E-Cadherin levels diminish as they travel in the mesodermal wings. Snail is a major EMT master transcription factor implicated in E-Cadherin down-regulation; it is critical for the adequate establishment of the three germ layers^6^. Interestingly, PS cells can delaminate in *Snail* knockout embryos, but nascent mesoderm maintains an intermediate level of E-cadherin and fails to migrate away^3^. PS cells asynchronously delaminate through apical constriction followed by retraction of the apical process and delamination on the basal side^4,7^. 3D time-lapse imaging of gastrulating mouse embryos revealed that apical constriction occurs in a pulsed ratchet-like fashion^8^.

In mouse, cell number is tightly regulated at the gastrulation stage. Indeed, experiments involving reduction or increase of embryo size highlighted a remarkable ability to compensate through change in proliferation rate^9,10^. Consistent with what was first observed by Snow in 1977^11^, immunostaining on sections and live imaging of mosaically labeled mouse embryos at the gastrulation stage showed that the frequency of mitoses is two times higher at the streak, compared to the rest of the epiblast^12^. Accordingly, single-cell RNA sequencing revealed that cells at the PS have a higher G2/M phase score than other epiblast cells^13^. Quantification of cell cycle length in rat embryos also indicated that cells at the PS are cycling faster than in the rest of the epiblast^14^.

The epiblast is a pseudostratified epithelium, where nuclei undergo Interkinetic Nuclei Migration (INM). In S phase, nuclei are basal. They are translocated toward the apical side during G2, so that mitosis occurs apically^15^. Live imaging confirmed the existence of such nuclear movements in the mouse epiblast^16^. Staining for Phospho-Histone H3, a marker for M phase, indicated that mitosis is indeed apical in the epiblast, except in the PS where about 40% of mitoses happen away from the apical pole^11,12^. Live imaging of embryos in which a small number of epiblast cells were labeled with membrane targeted GFP (thereby allowing tracking of cell shape and fate) showed that most cells undergoing non-apical mitotic rounding retained a basal attachment until cytokinesis, yet preferentially gave rise to one or two cells that delaminated to become mesoderm^12^. Non-apical mitoses have been described in other pseudostratified epithelia. For example, in the dorsal neuroepithelium of chick embryos, in which neural crest cells arise by EMT, non-apical mitoses are found in similar proportions than in the mouse PS. Importantly, abrogation of non-apical mitosis leads to a severe delay in neural crest cell delamination^17,18^.

All these observations indicate that PS cells have specific cycling properties and a high rate of non-apical mitoses that might be linked to their ability to extrude. We used experimental and in silico approaches to assess the functional relationship between proliferation rate, INM, cell contractility and the occurrence of non-apical mitoses in the epiblast and whether some of these parameters might actively contribute to cell delamination at the streak. Here we found that non-apical mitoses can be favored by multiple stimuli such as cytoplasmic actomyosin relaxation, a higher proliferation rate, a shorter G2, or detachment from the basement membrane. All these inputs act in concert at the PS. Importantly, non-apical mitoses promote but are not sufficient to provoke actual extrusion, which can only occur if tissue integrity is impaired.

## RESULTS

We previously established that non-apical mitoses at the PS primarily give rise to basal delamination of nascent mesoderm cells^12^. Therefore, we set out to identify the factors controlling the occurrence of non-apical divisions and assess whether they are causally linked to cell delamination. Variations in the position within the epithelium at which a cell divides could come from differences in apicobasal polarity, actomyosin contractility or cell cycle parameters (such as total duration of the cycle or duration of the G2/M phase). Thus, we performed a pharmacological screen targeting these cellular characteristics. While genetic manipulation allows for spatial control, it lacks precision in intensity and duration, which can complicate the interpretation of a phenotype, particularly in a highly dynamic context. The pharmacological approach addresses these limitations as it allows establishing optimal posology and recording the consequences of short-term alterations. One caveat is that all cells (epiblast and PS) are simultaneously targeted. To circumvent this, we used a computational model of proliferating pseudostratified epithelium in which all parameters can be controlled at the single cell level.

For biological experiments, we established a systematic protocol of measurements on embryos sectioned along the proximal-distal axis (Figure 1A, B). Sections were positioned along the anterior-posterior axis based on anatomical landmarks such as the long axis of the oval embryo shape and the morphology of visceral endoderm cells, as well as molecular markers including collagen IV for the basement membrane. The epiblast was divided into three regions: anterior and posterior halves of the epiblast, and PS, defined as the subregion of the posterior epiblast where the basement membrane is discontinuous. Staining for mitosis (Phospho-Histone H3) and nuclei (DAPI) allowed calculating the mitotic and non-apical mitotic indexes (Figure 1B). We quantified the percentage of basement membrane (collagen IV) degradation as a readout for EMT initiation and progression (Figure 1A).

**Figure 1:**
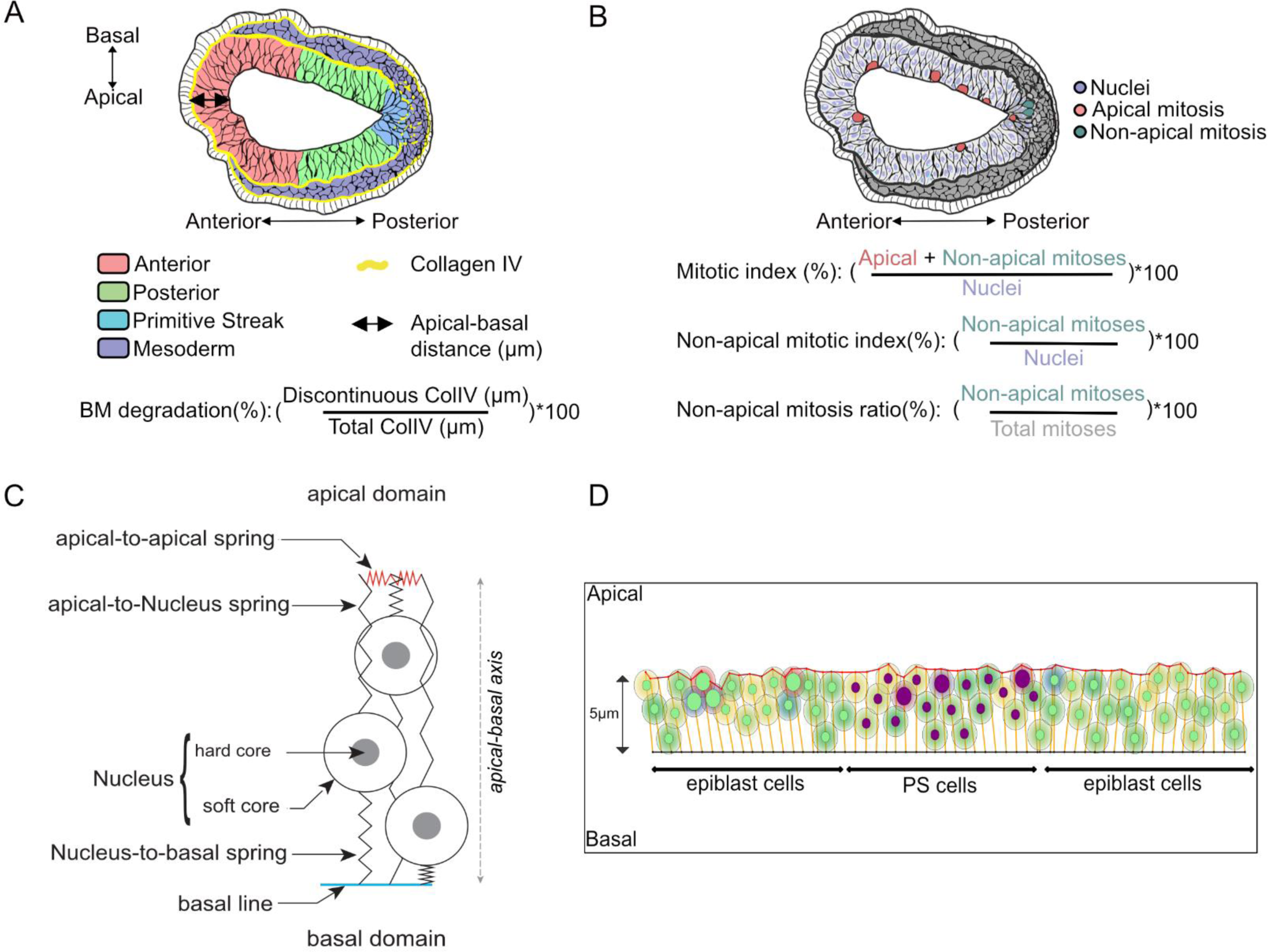
Calibration of pharmacological screening and in silico modeling. (A) Graphic representation of a transverse section from a E6.5 embryo. The epiblast is divided in three regions: anterior (pink), posterior (green) and the PS (PS, blue). Anterior and posterior regions are defined by dividing the epiblast in halves. The PS region is the subsection of the posterior region where the basement membrane (BM) is discontinuous (Collagen IV, yellow). Mesoderm cells that exited the epiblast are in purple. The black double head arrow represents the distance in µm between the apical and the basal surface. (B) Graphic representation of a transverse section from a E6.5 embryo. Nuclei are in purple, apical mitosis in pink and non-apical mitosis in green. A mitosis is considered non-apical if it occurs at least 10µm away from the apical surface (lumen), in a cell that retains apical and basal contact. (C) Diagram representing simulated cells controlled by various springs: apical cell adhesions (apical-to-apical spring), and the cytoplasmic cytoskeleton with basal cell-matrix adhesion (nucleus-to-basal spring and apical-to-Nucleus spring). Cell-matrix adhesions are modeled by the attachment of the nucleus-to-basal spring to the basal line. Nuclei are modeled with two layers. The soft layer can adjust to the environment to model pseudo-stratified morphology, the hard core cannot be deformed and increases during mitosis (D) Representation of a simulation setup (t=0h) for the in silico model adapted to the epiblast dynamics, with 19 PS (purple nuclei) cells and 40 epiblast cells (green nuclei). The red line represents the apical surface, and the black line the basal surface.

For the modeling of pseudo-stratified epithelium, we relied on a model originally developed on the chick neuroepithelium^19^. We used an extended version that allows the control of each parameter in time and space at the single cell level, described in ^20^. Briefly, each cell is approximated by a nucleus, an apical point, and a basal point. The two points are connected to the nucleus through dynamically adjustable springs representing the viscoelastic properties of the cytoplasm (Figure 1C). The nucleus is composed of an outer soft core and an inner hard core. Hard cores of neighboring cells cannot overlap while soft cores can but are subjected to a repulsion force. This allows representing packing and compression occurring between cells in a dense tissue. Apical cell-cell adhesions are modeled by springs between the apical points of neighboring cells (aa). On the basal side, basal points are attached to a fixed basal line representing the basement membrane. Basal points cannot overlap or swap positions. The cell cycle and INM movements are divided into three phases. The first phase corresponds to G1, S, and early G2, where movements are controlled by environmental constraints generated by neighboring cells. The second phase corresponds to late G2, where the pre-mitotic rapid apical movement (or PRAM) is simulated by contraction of the apical-nuclei string (aN) to bring nuclei to the apical pole, and extension of the basal-nuclei spring (bN) to accompany the movement. These settings are maintained in the third phase, the M phase, which results in the formation of two daughter cells. As the model is in 2D, most daughter cells are generated out of the 2D plane. To adapt the model to the epiblast size and configuration, a maximum height was set to allow no more than three pseudo-layers of nuclei, and dividing cells kept a basal attachment (Figure 1D).

### The difference in frequency and location of mitosis at the PS relative to the rest of the epiblast is independent of basement membrane degradation

In the mouse embryo, perforations in the basement membrane separating epiblast and visceral endoderm are distributed uniformly prior to migration of the Anterior Visceral Endoderm (AVE), the organizer that determines the anterior-posterior axis and therefore the position of the streak. Upon axis specification, perforations become more abundant in the posterior half of the embryo. These holes in the basement membrane are generated by a Nodal-dependent increase in the expression of Matrix Metalloproteinases (MMP) 2 and 14, and are necessary for embryo growth, PS morphogenesis, and gastrulation^21^. In EMT, loss of basal attachment due to basement membrane degradation is frequently associated with a disruption in apical-basal polarity. However, during mouse gastrulation, polarity markers are maintained in the PS^4,8,22^. This suggests that basement membrane degradation might be required for the occurrence of non-apical mitosis, independently of the loss of polarity.

To test this, we cultured pre-gastrulation E6 embryos for 6 hours with a combination of 20 µM Pan-MMPs inhibitor prinomastat hydrochloride and 100 µM MMP14 inhibitor, as previously described^21^. After incubation, embryos were fixed, and transverse cryosections were immunostained for Phospho-Histone H3, Collagen IV, nuclei (DAPI), and F-actin (phalloidin). Additionally, adjacent sections were stained for Cerberus 1 (Cer1) to identify the location of the AVE. We confirmed the abrogation of basement membrane degradation in embryos treated with MMPs inhibitors (Figure 2A, B). However, there was no significant difference in mitotic and non-apical mitotic indexes (Figure 2C, D), nor in the non-apical mitosis ratio (Supplementary 1A) throughout the epiblast relative to untreated embryos. After 8h of incubation (Supplementary 1D), some cells in the streak region acquired a mesenchymal morphology, but they were not able to cross the basement membrane (Figure 2E).

**Figure 2:**
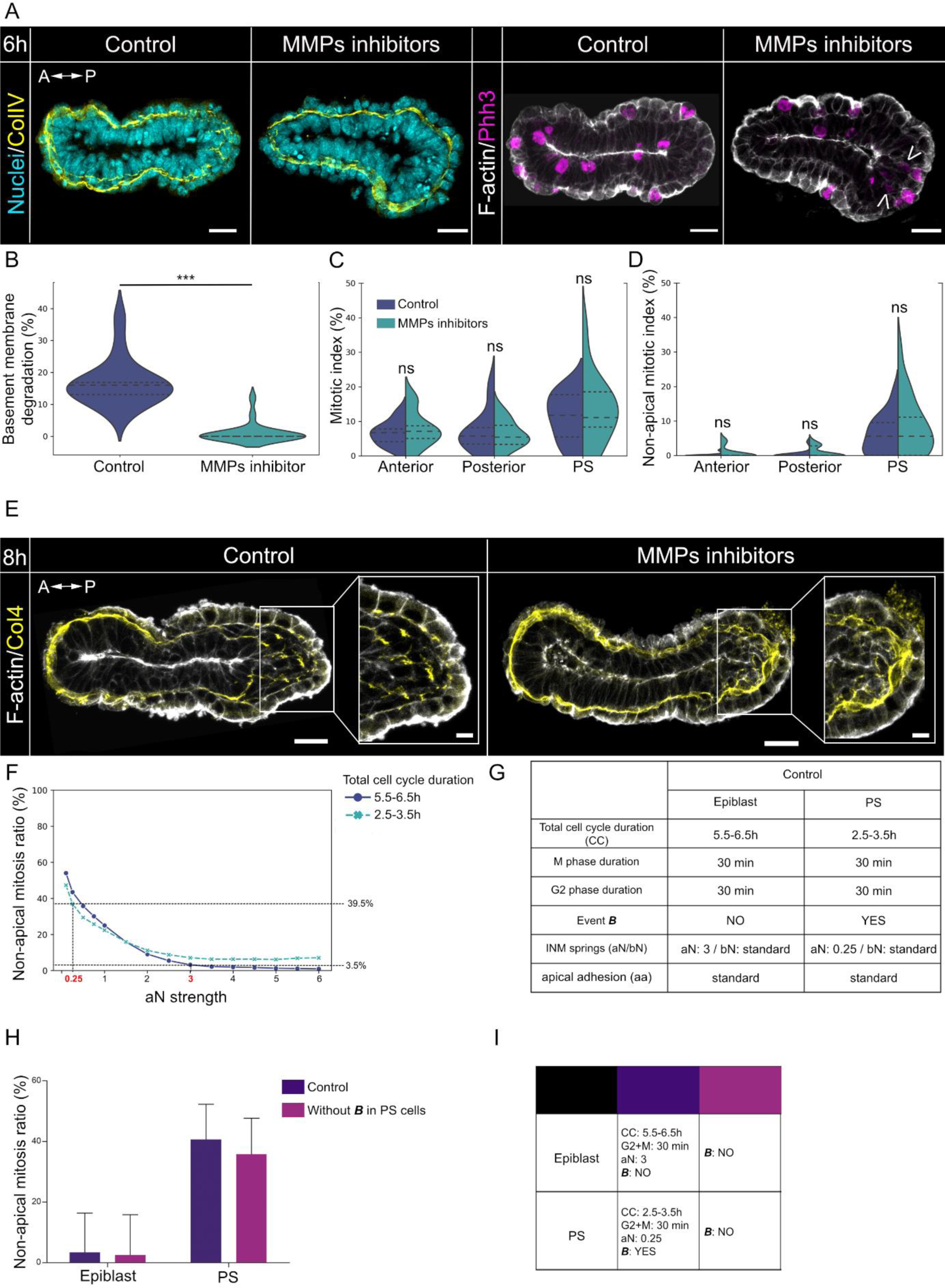
Preventing basement membrane degradation does not affect mitosis position. (A) Z-projection of transverse sections from E6 embryos cultured for 6 hours with control vehicle or MMPs inhibitors, stained for nuclei (DAPI, cyan), basement membrane (collagen IV, yellow), mitosis (Phospho-histone H3 (Phh3), magenta) and F-actin (phalloidin, grey). Arrow heads indicates instances of non-apical mitosis. Scale bars: 25µm. (B) Violin plots representing the percentage of basement membrane degradation in embryos cultured with control vehicle or MMPs inhibitors. Inner doted lines indicate third quartile, median, and first quartile. ***: P-value≤0.001. Normality was assessed using a Shapiro–Wilk test followed by Mann Whitney test. Controls: *n*= 14 slides from 8 embryos; MMPs inhibitors: *n*= 21 slides from 7 embryos. (C-D) Split violin plots representing the mitotic index (C) and non-apical mitotic index (D), in anterior, posterior and PS regions embryos cultured with control vehicle or MMPs inhibitors. Ns: nonsignificant. Normality was assessed using a Shapiro–Wilk test followed by Mann-Whitney or t-test. Controls: *n*= 14 slides from 8 embryos, anterior: 694 nuclei, posterior: 436 nuclei, PS: 332 nuclei. MMPs inhibitors: *n*= 21 slides from 7 embryos, anterior: 846 nuclei, posterior: 660 nuclei, PS: 441 nuclei. (E) Z-projection of transverse sections from E6 embryos cultured for 8 hours with control vehicle or MMPs inhibitors, stained for basement membrane (collagen IV, yellow) and F-actin (phalloidin, grey). White squares show regions of zooms in the PS regions. Scale bars: 25 µm and 10 µm (zooms). (F) Graph from in silico simulations representing the percentage of non-apical mitosis depending on aN strength in epiblast cells (6h cell cycle) and PS cells (3h cell cycle). 200 simulations with 30 proliferating cells. (G) Table presenting the simulated parameters set for epiblast and PS cells in control conditions in the in silico epiblast model. Event B designate the detachment of cells’ basal point from the basal line. (H) Bar plot representing the ratio of non-apical mitosis (%), in the region of the epiblast and PS in simulations ran with control settings or in situations where PS cells do not undergo event ***B***. 200 simulations with 20 epiblast cells and 19 PS cells. (I) Table recapitulating the settings of epiblast and PS cells in control and test simulations. For the simulation, only the modified parameters are indicated. A-P: Anterior – Posterior; Ap-Bas: Apical – Basal; PS: PS; CC: cell cycle; aN: apical-Nuclei spring, bN: basal-Nuclei spring; aa: apical adhesion spring.

Before modelling the impact of detachment from the basement membrane in silico, we first had to calibrate the parameters of cell cycle and INM allowing to recapitulate the percentage of non-apical mitoses observed in vivo. As the velocity of PRAM is controlled by the contraction of the apical-to-Nucleus (aN) spring in the model, we tested a range of strength values for aN (Figure 2F). We simulated 200 repetitions of 30 proliferating cells with a cell cycle of 6 ± 0.5h or 3 ± 0.5h, corresponding to measured values for epiblast and PS cells, respectively^11,14^. The percentages of non-apical mitoses observed at the PS (circa 40%) could only be reached at very low values of aN, suggesting that an almost complete loss of PRAM is necessary. In addition, we noted that a faster cell cycle favors non-apical mitoses even at higher PRAM intensity. Indeed, the frequency of non-apical mitoses for medium values of aN reached a plateau at 7% with a cell cycle length of 3h, and only 1% with a length of 6h. Based on these simulations, we used the following as default parameters: 6 ± 0.5h of cell cycle and 3 of aN strength for epiblast cells, 3 ± 0.5h of cell cycle and 0.25 of aN strength for PS cells (Figure 2G). We counted the number of PS cells on cross sections of stage E6.5 mouse embryo and found a mean population size of 19 PS cells per section^12^. Thus, we set the size of the PS population at 19 cells, surrounded by 20 epiblast cells on each side. Simulations start with a period of 6 hours during which all cells are attached basally. Then there is an 18h long window of opportunity for PS cells to detach from the basal line, which is referred as event ***B*** (Figure 2G, F) and represents basement membrane degradation^20^.

We then ran simulations with or without detachment of PS cells from the basal line and plotted the rate of non-apical mitoses (Figure 2H, I). As observed in embryos treated for 6h with MMPs inhibitors, impairing detachment from the basal line did not suppress the occurrence of non-apical mitoses at the PS (Figure 2I, J).

Collectively, these data indicate that preventing basement membrane degradation does not influence the rate of mitoses and non-apical mitoses at the primitive streak.

### Actomyosin relaxation is sufficient to drive the appearance of non-apical mitosis in the epiblast

As INM movements are usually driven by actomyosin in short pseudo-stratified epithelia^15^, we tested whether a regional modification in actomyosin contractility could be responsible for the increased rate of non-apical mitosis at the PS.

The shape of nuclei in interphase can be used as a readout of the mechanical constraints exerted on cells^23,24^, which are believed to be particularly high in pseudo-stratified epithelia^25,26^. We marked nuclear membrane via staining for Lamin B1, the first Lamin subtype expressed during early development in mouse^27,28^, and calculated the nuclei size ratio by dividing the short axis length by the long axis length (Figure 3A). A ratio close to 0 corresponds to an elongated nucleus, indicating that the cytoskeleton is contracted. A ratio close to 1 indicates a round nucleus, reflecting relaxation of the cytoskeleton network. Nuclei shape ratios were recorded in the anterior, posterior, and PS regions of E6.25, E6.5, and E7 embryos (Figure 3B, B”). Nuclei in the PS were rounder, compared to the rest of the epiblast, from E6.25 onwards, with a peak at E6.5 and maintenance until E7 (Figure 3C-C’’). This is compatible with a local relaxation of the cytoskeleton at the streak, leading to nuclei shape changes, upon the onset of gastrulation.

**Figure 3:**
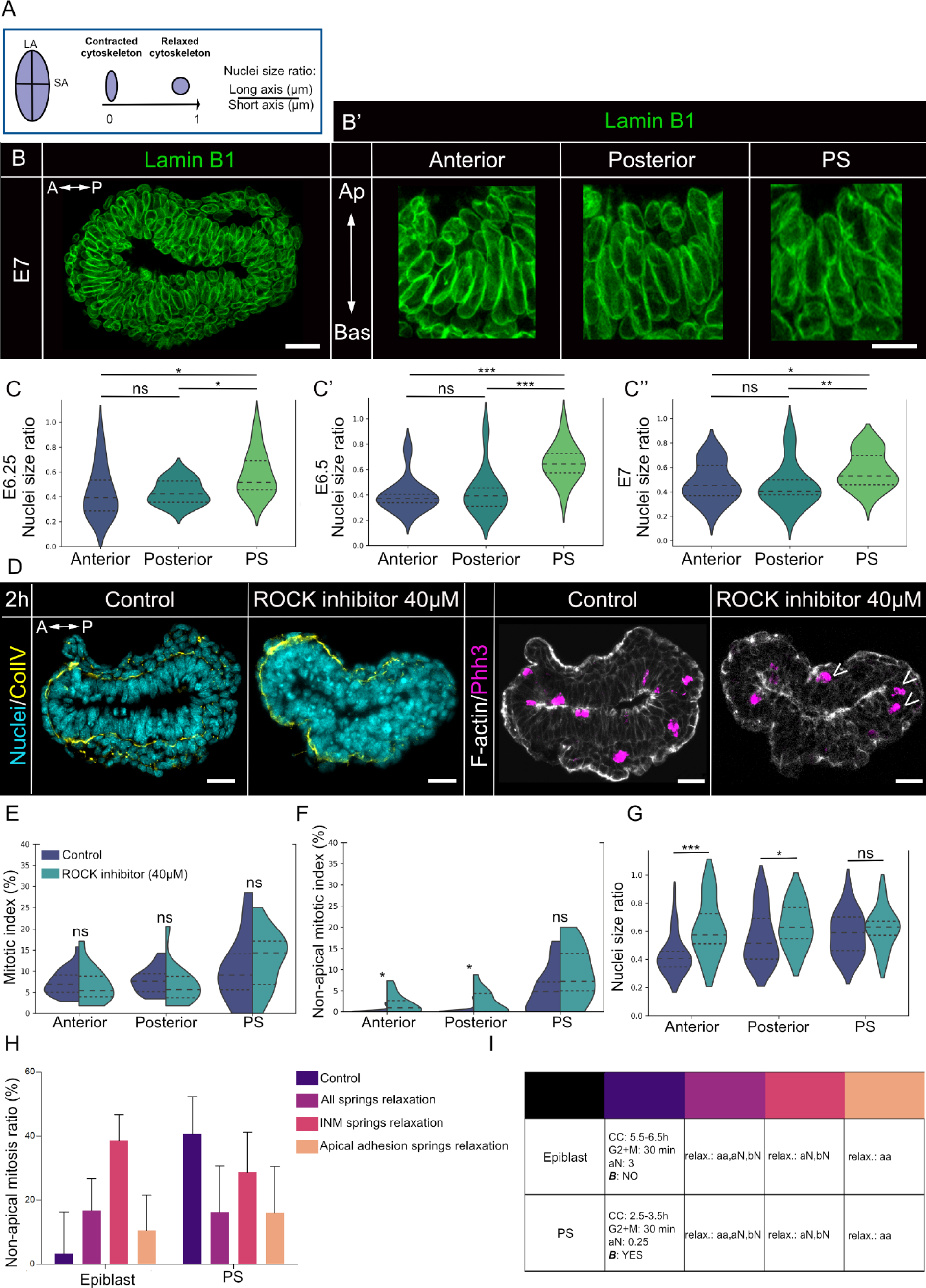
Actomyosin relaxation at the PS triggers the disharmonization of cell cycle and pre-mitotic rapid apical movements. (A) Graphic representation of a nucleus. Nuclei size ratio is calculated by dividing the short axis (SA) by the long axis (LA) of each nucleus. A ratio close to 0 indicates an elongated shape, while a ratio close to one indicates a round shape. (B-B’) Z-projection of a transverse section from a E7 embryo, stained for Lamin B1 (green) (B). Zoom on nuclei from the anterior, posterior and PS regions (B’). Scale bars: 25µm (D) and 10µm (B’). (C-C”) Violin plots representing nuclei size ratio at the anterior, posterior and PS region at E6.25 (C), E6.5 (C’) and E7 (C”). Inner doted lines indicate third quartile, median, and first quartile. Normality was assessed using a Shapiro–Wilk test followed by a Kruskall-Wallis test with Dunnet post hoc. ns: non-significant, *: P-value≤0.05, **: P-value≤0.01and ***: P-value≤0.001. E6.25: *n*= 4 embryos, anterior: 20 nuclei, posterior: 20 nuclei, PS: 20 nuclei. E6.5: *n*= 4 embryos, anterior: 20 nuclei, posterior: 20 nuclei, PS: 20 nuclei. E7: *n*= 4 embryos, anterior: 20 nuclei, posterior: 20 nuclei, PS: 21 nuclei. (D) Z-projection of transverse sections from E6.5 embryos cultured for 2 hours with control vehicle or ROCK inhibitor at 40µM, stained for nuclei (DAPI, cyan), basement membrane (collagen IV, yellow), mitosis (Phh3, magenta) and F-actin (phalloidin, grey). Arrow heads indicates instances of non-apical mitoses. Scale bars: 25µm. (E-F) Split violin plots representing the mitotic index (E) and non-apical mitotic index (F), in anterior, posterior, and PS regions in embryos cultured with control vehicle or 40µM ROCK inhibitor. Inner doted lines indicate third quartile, median, and first quartile. Ns: nonsignificant, *: P-value≤0.05. Normality was assessed using a Shapiro–Wilk test followed by Mann-Whitney or t-test. Control: *n*= 26 slides from 12 embryos, anterior: 1048 nuclei, posterior: 663 nuclei, PS: 348 nuclei. ROCK inhibitor: *n*= 14 slides from 7 embryos, anterior: 738 nuclei, posterior: 531 nuclei, PS: 205 nuclei. (G) Violin plots representing nuclei size ratio in anterior, posterior, and PS regions, in embryos cultured with control vehicle or 40µM ROCK inhibitor. Inner doted lines indicate third quartile, median, and first quartile. Ns: non-significant, *: P-value≤0.05, ***: P-value≤0.001. Normality was assessed using a Shapiro– Wilk test followed by Mann-Whitney or t-test. Control: *n*= 11 embryos, anterior: 37 nuclei, posterior: 37 nuclei, PS: 37 nuclei. 40µM ROCK inhibitor: *n*= 5 embryos, anterior: 19 nuclei, posterior: 20 nuclei, PS: 20 nuclei. (H) Bar plot representing the percentage of non-apical mitosis in the region of the epiblast and PS in various simulations: control, relaxation of all springs, relaxation of INM (aN/bN) springs, and relaxation of apical adhesion springs. 200 simulations with 20 epiblast cells and 19 PS cells. (I) Table recapitulating the settings of epiblast and PS cells in control and test simulations. For the simulation, only the modified parameters are indicated. A-P: Anterior – Posterior; Ap-Bas: Apical – Basal; PS: primitive streak; INM: interkinetic nuclei migration; CC: cell cycle; ***B***: detachment of basal adhesions from the basal line; aN: apical-Nuclei spring, bN: basal-Nuclei spring; aa: apical adhesion spring.

The small Rho-GTPase RhoA regulates both actomyosin contractility and cell cycle progression. In addition, during EMT, RhoA participates in the cellular changes required to acquire a mesenchymal-like phenotype and migrate^29^. Rho-associated kinase (ROCK), a downstream effector of RhoA, regulates actomyosin contractility via myosin light chain phosphorylation^30,31^. In the pseudostratified epithelium of the *Drosophila* imaginal discs, pharmacological treatment with ROCK inhibitor Y27632 leads to basal accumulation of nuclei and the appearance of non-apical mitoses^32^. To investigate the link between mitosis location and actomyosin dynamics in the mouse epiblast, we cultured E6.5 embryos for 2h with 40 µM Y27632. Embryos were then fixed, and transverse cryosections were immunostained for Phospho-Histone H3, Collagen IV, nuclei (DAPI), and F-actin (phalloidin). There was no change in the mitotic index (Figure 3D, E). However, a 2-fold increase in the non-apical mitotic index (and a 10-fold increase of the non-apical mitosis ratio) were recorded in the anterior and posterior epiblast (Figure 3D, F, Supplementary 1B). This is compatible with an actomyosin-driven INM in the epiblast: indeed, it is likely that disturbing actomyosin contractility affects the ability of cells to drive their nuclei to the apical surface in G2, which results in mitosis occurring in ectopic location. Interestingly, non-apical mitotic index remained unchanged at the PS (Figure 3D, F), strongly suggesting that actomyosin contractility is intrinsically low in PS cells. Upon ROCK inhibition, nuclei shape was rounder in the anterior and posterior epiblast (Figure 3G), confirming it induced a relaxation of the actomyosin cytoskeleton, while the shape of nuclei in the PS did not change (Figure 3G). The effect of the inhibitor was reversible, as embryos treated for 2h with 40 µM of ROCK inhibitors then washed were able to adjust rapidly and survive overnight in culture (Supplementary 1E).

To further explore the influence of cytoskeleton contractility on the occurrence of non-apical mitoses, we performed simulations with a reduced strength for all springs: aN and bN for the cell body/cytoplasm and aa for apical cell-cell adhesion (Figure 3H, I; light purple). Interestingly, such global relaxation increased the frequency of non-apical mitoses in the epiblast and reduced it in the PS (Figure 3H). Thus, we set out to assess the relative contributions of the cytoplasmic springs (aN, bN) that control INM/PRAM and of the apical adhesion spring (aa), by relaxing them independently (Figure 3H, I; pink and orange). Relaxing aN/bN strongly promoted non-apical mitoses in the epiblast but did not affect PS cells. Relaxing aa mildly promoted non-apical mitoses in the epiblast and lowered their frequency in the PS. It is interesting to note that in all simulations with aa relaxation the tissue experienced severe deformations (Figure 3H, I). The changes obtained in the scenario where all springs were relaxed are compatible with what we observed in embryos treated with a high dose of ROCK inhibitor (400 µM): those displayed severe morphological defects (Supplementary 2A, B); the mitotic index was decreased in all regions of the epiblast (Supplementary 2C) while the non-apical mitotic index was increased in the anterior and posterior epiblast but decreased in the PS (Supplementary 2D). We thus interpret that the effects observed at 40 µM of ROCK inhibitor correspond to a mild inhibition of the cell cytoskeleton contractility that can be modeled by the relaxation of only the INM springs.

Collectively, these experimental and simulated data indicate that a mild inhibition of ROCK generates non-apical mitoses in the epiblast primarily by affecting cytoplasmic actomyosin activity rather than apical actomyosin. This suggests that INM is governed by actomyosin in the epiblast and that, comparatively, actomyosin contractility is weak in PS cells, leading to the appearance of non-apical mitoses.

### Cell cycle regulation is less stringent in the PS, compared to the rest of the epiblast

The complex formed by Cyclin-dependent Kinase 1 (CDK1) and CyclinD1 is critical for the control of cell entry into mitosis. In the avian neuroepithelium, overexpression of CyclinD1 leads to the appearance of non-apical mitoses^33^. In zebrafish retina, CDK1 is required for the rapid basal to apical movement of nuclei during G2^34^. Therefore, we chose to pharmacologically target CDK1, negatively using a CDK1 inhibitor (R03366), and positively via a Wee1 inhibitor (PD16685). Wee 1 is a nuclear kinase that regulates mitosis by inhibiting CDK1 through phosphorylation. Wee1 inhibition enables to bypass the G2/M checkpoint^35^.

A 2h incubation with 175 µM of CDK1 inhibitor abrogated all mitosis, confirming that targeting CDK1 regulation can rapidly affect the cell cycle in the epiblast (Figure 4A-C). Embryos were able to recover from the treatment and developed normally (Supplementary 1E). With the goal to increase the frequency of mitoses without major morphological anomaly, we performed a dose-response analysis for the Wee1 inhibitor (5, 10, 20, 40, and 80 µM for 2 hours). Embryos were fixed directly after incubation. We measured the percentage of basement membrane degradation as well as the mitotic and non-apical mitotic indexes on transverse cryosections immunostained for Phospho-Histone H3, Collagen IV, nuclei, and F-actin (Figure 4D). In the anterior and posterior epiblast, we observed a dose-dependent increase of the mitotic index, indicating an acceleration of the cell cycle, as well as an increase of the non-apical mitotic index and non-apical mitosis ratio (Figure 4D-F; Supplementary 1C). We previously showed that an acceleration of the cell cycle is sufficient to trigger non-apical mitoses (Figure 2F), therefore some of the increase of non-apical mitoses in the epiblast upon treatment with Wee1 inhibitor might be linked to the global increase in mitotic index. However, as PRAM take place in G2, Wee1 inhibition might also contribute to the higher rate of non-apical mitoses by shortening the window of opportunity for PRAM to occur. By contrast, the frequency and location of mitoses in the PS stayed constant except at high doses of drug (40-80 µM) at which most cells were entering mitosis (Figure 4E, F). These data indicate that cells at the PS were less sensitive to Wee1 inhibition compared to the rest of the epiblast, strongly suggesting an intrinsically shorter cell cycle and G2 duration in PS cells.

**Figure 4:**
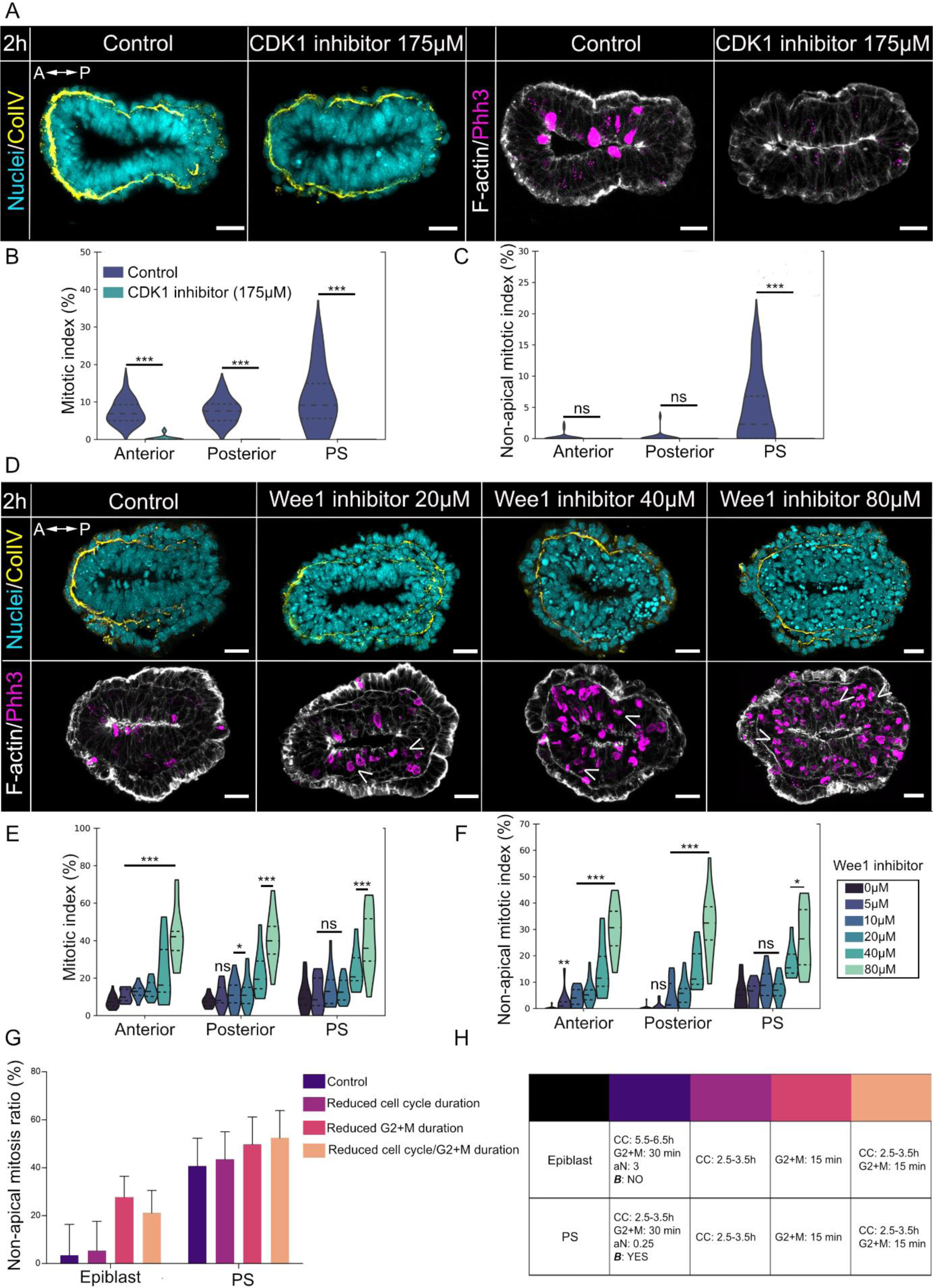
Cell cycle regulation at the PS is less stringent than in the rest of the epiblast, which triggers the appearance of non-apical mitoses. (A) Z-projections of transverse sections from E6.5 embryos cultured for 2 hours with control vehicle or 175µM of CDK1 inhibitor (R03366), stained for nuclei (DAPI, cyan), basement membrane (collagen IV, yellow), mitosis (phh3, magenta) and F-actin (phalloidin, grey). Scale bars: 25µm. (B-C) Violin plots representing the mitotic index (B) and non-apical mitotic index (C) in anterior, posterior, and PS regions in embryos cultured with control vehicle or 175µM CDK1 inhibitor. Inner doted lines indicate third quartile, median, and first quartile. ***: P-value≤0.001. Normality was assessed using a Shapiro–Wilk test followed by Mann-Whitney test. Control: *n*= 24 slides from 8 embryos, anterior: 923 nuclei, posterior: 595 nuclei, PS: 312 nuclei. 175µM CDK1 inhibitor: *n*= 11 slides from 7 embryos, anterior: 414 nuclei, posterior: 329 nuclei, PS: 124 nuclei. (D) Z-projection of transverse sections from E6.5 embryos cultured for 2 hours in vehicle (Control) or with 20µM, 40µM or 80µM of Wee1 inhibitor, stained for nuclei (DAPI, cyan), basement membrane (collagen IV, yellow), mitosis (phh3, magenta) and F-actin (phalloidin, grey). Arrow heads indicates instances of non-apical mitoses. Scale bars: 25µm. (E-F) Violin plots representing the mitotic index (E) and non-apical mitotic index (F), in anterior, posterior, and PS regions of embryos cultured with control vehicle or 5µM, 10µM, 20µM, 40µM or 80µM of Wee1 inhibitor for 2 hours. Inner doted lines indicate third quartile, median, and first quartile. ns: nonsignificant, *: P-value≤0.05, **: P-value≤0.01 and ***: P-value≤0.001. Normality was assessed using a Shapiro–Wilk test followed by Mann-Whitney or t-test. Control: *n*= 27 slides from 13 embryos, anterior: 1111 nuclei, posterior: 705 nuclei, PS: 362 nuclei. 5µM Wee1 inhibitor: *n*= 12 slides from 5 embryos, anterior: 501 nuclei, posterior: 298 nuclei, PS: 166 nuclei. 10µM Wee1 inhibitor: *n*= 16 slides from 10 embryos, anterior: 647 nuclei, posterior: 412 nuclei, PS: 324 nuclei. 20µM Wee1 inhibitor: *n*= 21 slides from 9 embryos, anterior: 1236 nuclei, posterior: 716 nuclei, PS: 555 nuclei. 40µM Wee1 inhibitor: *n*= 9 slides from 4 embryos, anterior: 409 nuclei, posterior: 262 nuclei, PS: 226 nuclei. 80µM Wee1 inhibitor: *n*= 12 slides from 6 embryos, anterior: 575 nuclei, posterior: 330 nuclei, PS: 297 nuclei. (G) Bar plot representing the percentage of non-apical mitosis in the region of the epiblast and PS in various simulation: control, reduction of the total cell cycle duration (3h), reduction of G2 and M phase duration (15 minutes) and reduction of both the total duration of the cell cycle (3h) and of G2 and M duration (15 minutes). 200 simulations with 20 epiblast cells and 19 PS cells. (H) Table recapitulating the settings of epiblast and PS cells in control and test simulations. For the simulation, only the modified parameters are indicated. A-P: Anterior – Posterior; Ap-Bas: Apical – Basal; PS: primitive streak; CC: cell cycle; *B*: detachment of basal adhesions from the basal line; aN: apical-Nuclei spring, bN: basal-Nuclei spring; aa: apical adhesion spring.

To assess the relative impact of G2 and total cell cycle duration, we performed simulations in which we modulated their duration independently or simultaneously. The model has a simplified version of the cell cycle in which PRAM forces are applied through G2 and M. Thus, to mimic the effect of the Wee1 inhibitor in shortening biological G2 and the associated duration of PRAM forces, we reduced the total duration of simulated G2+M by half. In the control situation, epiblast cells cycle every 6 ± 0.5h and PS cells every 3 ± 0.5h. All cells have a G2/M of 30 minutes each (Figure 4G, H; dark purple). If all cells were forced to cycle every 3 ± 0.5h at constant G2/M duration, it only generated a marginal increase of non-apical mitoses in the epiblast (Figure 4G, H; light purple). If both cell types kept their respective cell cycle length but were forced to have a G2/M of 15 minutes each, there was a massive increase of non-apical mitoses in the epiblast (Figure 4G, H; pink). If both changes were combined (cell cycle 3 ± 0.5h, G2/M 15 min in all cells), it did not generate a further increase of non-apical mitoses in the epiblast (Figure 4G, H; orange). By contrast, all these changes had negligible effects on the PS. These results indicate that specifically affecting G2/M duration has more impact on the rate of non-apical mitoses than shortening the whole cell cycle without affecting G2/M duration. Overall, these data indicate that an acceleration of the cell cycle and a shortened G2 duration contribute to the increased incidence of mitoses and non-apical mitoses observed at the primitive streak.

### The rate of non-apical mitoses correlates with the occurrence of basal extrusion

In view of the results from pharmacological screening and in silico modeling, we conclude that relaxation of cytoplasmic actomyosin, a faster cell cycle, and a shortening of the G2 phase collectively contribute to the increased incidence of non-apical mitoses at the PS during gastrulation. Given that in vivo time-lapse imaging showed that cells undergoing non-apical mitosis primarily gave rise to basally extruding mesodermal cells^12^, we wondered if those mechanisms might also be upstream of basal extrusion.

Interestingly, computational modeling of EMT in a tall pseudostratified epithelium using the same model indicated that the position of nuclei correlates with the directionality of extrusion^20^. In particular, basal positioning (defined as more basal than the average position of nuclei) specifically favors basal extrusion. Therefore, some of the experimental and simulated experimental conditions we tested might have a broader impact on nuclei distribution in the tissue than the position of mitoses. However, in a comparatively flat epithelium like the gastrulating epiblast it is unclear whether basal positioning has any importance for extrusion given that all cells have their nuclei relatively close to the basement membrane at any time point.

To assess this, we plotted the rate of basal positioning of nuclei in epiblast and PS for all simulated conditions (Figure 5A-B). There was a high rate of basal positioning in control PS cells. Preventing detachment from the basal line or enforcing normal actomyosin contractility (Figure 5A, PS experimental conditions 2 to 5) prevented basal positioning of nuclei in PS cells. By contrast none of the treatments affecting contractility or cell cycle parameters were able to promote basal positioning in epiblast cells, where only a forced detachment from the basal line was sufficient to generate a high rate of basal positioning (Figure 5A, epiblast experimental condition 2).

**Figure 5:**
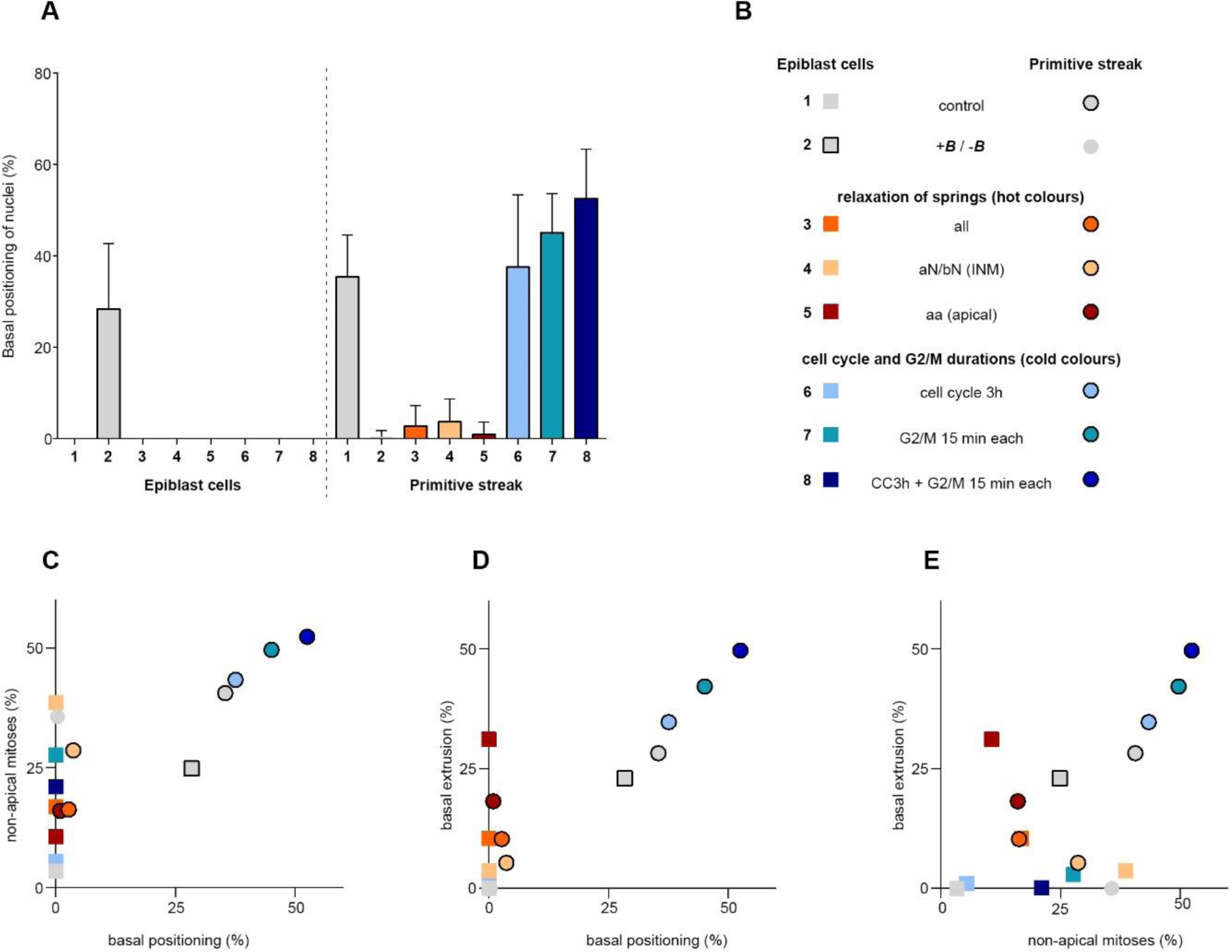
Non-apical mitoses are positively correlated with basal extrusion at the primitive streak. (A) Bar plot of the mean percentage of nuclei basal positioning in tested scenarios for epiblast and PS cells. (B) Legend for tested scenarios. Squares correspond to the region of the epiblast and circles the PS. (C) Scatter plot of the percentage of non-apical mitoses and mean rate of nuclei basal positioning in tested scenarios for the epiblast and PS. (D) Scatterplot of the mean rate of basal extrusion and basal positioning in tested scenarios for the epiblast and the PS. (E) Scatter plot of the mean rate of basal extrusion and the percentage of non-apical mitosis in tested scenarios for the epiblast and PS.

Next, we wondered if overall basal positioning of nuclei and the occurrence of non-apical mitoses were correlated (Figure 5C). This revealed two different patterns: in epiblast cells, non-apical mitoses occurred in absence of basal positioning unless loss of basal attachment was enforced (grey square with black outline). In PS cells both parameters were strongly correlated unless we imposed a relaxation of the springs throughout the tissue, which completely canceled basal positioning but only lowered the rate of non-apical mitoses (Figure 5C, circles in hot colors). These data indicate that non-apical mitoses and overall basal positioning of nuclei can be uncoupled. This means that non-apical mitoses can occur even in absence of a global redistribution of nuclei in the tissue.

Next, we looked at a putative correlation between basal positioning and basal extrusion (Figure 5D) and found a similar relationship: there was a strong positive correlation in PS cells unless contractility was affected (circles in hot colors), and no correlation in the epiblast unless epiblast cells were forced to detach from the basal line (grey square with a black outline). Finally, we looked at the relationship between non-apical mitoses and basal extrusion. Here, the data were not split in two groups, and we found an overall positive correlation between the two factors (Pearson r = 0.59, p = 0.0157).

These analyses revealed that non-apical mitoses can specifically occur in absence of overall tissue disorganization leading to massive basal positioning. In addition, non-apical mitoses are positively correlated with basal extrusion. However, it is important to note that extrusion itself only occurs significantly if cells detach from the basal line or if all springs or apical springs are relaxed. This means that actual delamination requires a condition in which epithelial integrity is impaired.

### The occurrence of non-apical mitoses is regulated independently of EMT progression

In control conditions, non-apical mitoses are followed by basal delamination of mesodermal precursors^12^. Thus, we wondered if any of the pharmacological treatments that led to ectopic non-apical mitoses might be sufficient to promote ectopic EMT in the epiblast. We monitored markers of EMT progression, notably the expression of the EMT marker Snail1, the pattern of E-cadherin and N-cadherin, and the percentage of basement membrane degradation, upon pharmacological inhibition of ROCK, Wee1, and CDK1.

In embryos treated with 40 µM of ROCK inhibitor, immunostaining for Snail1 showed that cells at the PS were still able to transition, and no ectopic EMT site was detected (Figure 6A). This was confirmed by staining for E-cadherin and N-cadherin, as there was no alteration in the cadherin switch (Figure 6B). Similarly, we observed no significant change regarding the percentage, pattern, or localization of basement membrane degradation (Figure 3D; Figure 6D).

**Figure 6:**
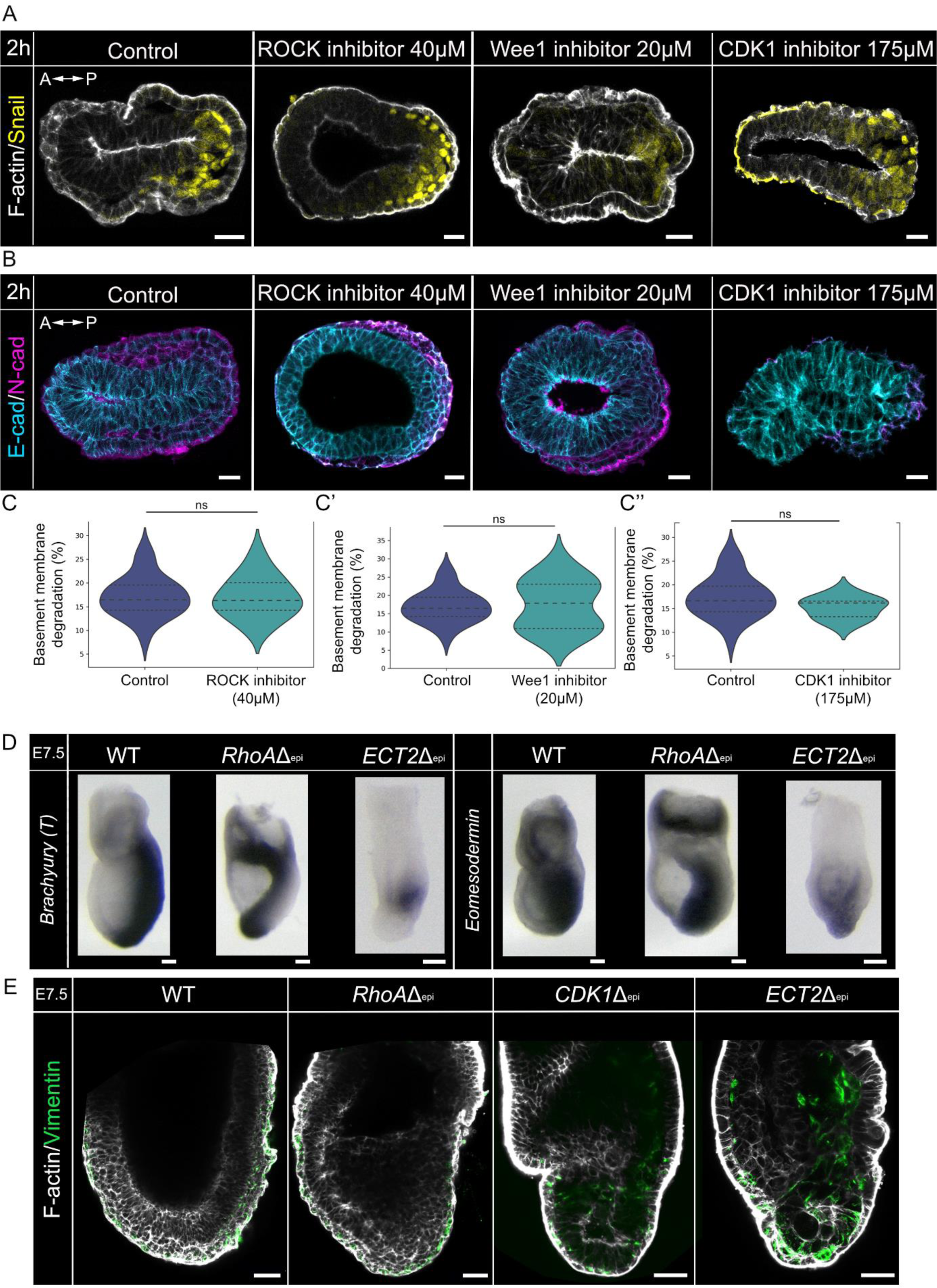
Dynamic changes observed at the PS are independent from epithelial-mesenchymal transition progress and mesoderm specification *in vivo*. (A-B) Z-projections of transverse sections from E6.5 embryos cultured for 2 hours with control vehicle, 40µM of ROCK inhibitor, 20µM Wee1 inhibitor or 175µM CDK1 inhibitor, stained for the EMT marker Snail (yellow) and F-actin (phalloidin, grey) (A) or E-cadherin (cyan) and N-cadherin (magenta) (B). Scale bars: 25µm. (D-D”) Violin plots representing the percentage of basement membrane degradation in embryos cultured with control vehicle, 40µM ROCK inhibitor (D), 20µM Wee1 inhibitor (D’), 175µM CDK1 inhibitor (D”). Control: *n=* 27 slides from 13 embryos; 40µM ROCK inhibitor: *n=* 14 slides from 7 embryos; 20µM Wee1 inhibitor: *n=* 21 slides from 9 embryos; 175µM CDK1 inhibitor: *n=* 11 slides from 7 embryos. (E) In situ hybridization for Brachyury (T) (top), and Eomesodermin (bottom) on E7.5 WT, RhoAΔepiblast, Ect2Δepiblast embryos. Scale bars: 100 µm. Brachyury (T): RhoAΔepiblast: *n=*2, ECT2Δepiblast: *n=*4, Eomesodermin: RhoAΔepiblast: *n=*5; Ect2Δepiblast: *n=*5.

For Wee1, we focused on embryos treated with 20 µM of the inhibitor for 2 hours. Immunostaining for Snail1, E-cadherin, and N-cadherin indicated that EMT was initiated normally at the PS (Figure 6A, B). While these conditions lead to a global increase of non-apical mitoses in all the epiblast (Figure 4D, F; Supplementary 1C), there were no ectopic site of EMT nor significant change regarding the percentage of basement membrane degradation (Figure 4D, Figure 6A, B, D’). In addition, preventing entry in mitosis by culturing embryos for 2 hours with 175 µM of CDK1 inhibitor did not affect basement membrane degradation, the expression of Snail1, or the cadherin switch (Figure 4A; Figure 6A, B, D”), indicating that cell division per se is not necessary for EMT initiation.

These data indicate that while local changes in actomyosin contractility and cell cycle parameters at the PS likely favor the pre-positioning of cells for exit, they are not sufficient to trigger or block EMT progression.

### Relationships between EMT, basal delamination, mesoderm specification, and mesoderm migration are complex and nonlinear

There is evidence that mitosis can influence gene regulation and EMT progression, notably through the process of gene bookmarking^36,37^. Therefore, we asked whether mitosis frequency and location could favor mesoderm specification.

We examined the phenotypes of embryos carrying epiblast-specific deletion of the small GTPase *RhoA*, the Guanine exchange factor *Ect2*, and the kinase *Cdk1*, which are involved in cytoskeleton regulation, cytokinesis, and cell cycle regulation, respectively. Δ^epiblast^ embryos were obtained from the combination of *RhoA, Ect2 and Cdk1* conditional alleles^38–40^ and a Cre-recombinase under the dependency of the *Sox2* promoter, which is turned on in the prospective epiblast at E3.5^41^ (Figure E, Supplementary 3).

Deletion of *RhoA* in the epiblast led to disruption of the pseudostratified arrangement, detectable from E6,5 onwards (Supplementary 3A-G). There appeared to be a trend towards a higher mitotic index at E6.5, possibly due to delays in cytokinesis (Supplementary 3C, D). Basement membrane degradation was detectable in the posterior region, and cells with a mesenchymal phenotype accumulated at the streak and into the lumen (Supplementary 3E). This is compatible with an adequate initiation of EMT but a defect in cell migration away from the PS, like what was observed in *Rac1* epiblast-specific mutants^22^, or *Rac1* and *RhoA* mesoderm-specific mutants^42^. By E7.5, *RhoA*Δ^epiblast^ mutants exhibited severe morphological defects. Lumens could be spotted within the accumulation of cells on the posterior side of the embryo, suggesting a disruption of polarity (Supplementary 3F). Similarly, epiblast-specific deletion of *Cdk1* or the cytokinesis regulator *Ect2* both resulted in a severe phenotype with developmental arrest (Figure 5E, Supplementary 3H, I). In *RhoA*Δ^epiblast^, as well as *CDK1*Δ^epiblast^ and *ECT2*Δ^epiblast^ mutants, mesoderm identity was specified, as shown by the expression of the PS and nascent mesoderm marker *Brachyury* (T) (Figure 6D), and Vimentin (Figure 6E). These data underscore the critical role for RhoA, CDK1 and ECT2 in maintaining epiblast integrity, basal delamination at the PS, and cell migration.

## DISCUSSION

At gastrulation, two novel germ layers are generated by delamination of cells from the epiblast. This process necessarily involves an increase in cell number at the site of cell exit that is superior to the one required for homogeneous embryo growth. In chick embryos, global movements in the epiblast precede gastrulation and position PS precursors along the antero-posterior axis^43^. The mouse PS, however, was shown to arise in situ, without large scale displacement of epiblast cells^4^. A higher rate of cell division at the streak therefore appears as a suitable mechanism to fuel gastrulation EMT^11,12^. In addition to its quantitative input, mitosis may also play a qualitative role, notably through the mechanical impact of mitotic rounding on epithelial architecture^44^. The apicobasal position of mitosis may be of importance, as one would instinctively postulate that cells dividing closer to the basal pole have an advantage for subsequent delamination, even if they only detach apically at the late steps of cytokinesis.

Neural crest delamination from the chick neuroepithelium is a well-studied example of EMT occurring in a middle size pseudostratified epithelium, where the impact of non-apical mitosis on cell exit was clearly demonstrated ^17^. There, biological and computational data highlighted a role for the reduction of INM in basal positioning and subsequent basal delamination^20^. Loss of INM in EMT cells increases the probability of basal positioning, while maintenance of INM in surrounding non EMT cells results in apical crowding and could mechanically favor their delamination^20^. Interestingly, we find a similar pattern in the epiblast, a short pseudostratified epithelium where all cells are relatively close to the basal pole. An additional point of similarity between those EMT events lies in the temporal regulation of INM. Indeed, non-apical mitoses were detected prior to the initiation of EMT in the pre-streak posterior epiblast^12^ and the prospective neural crest cells^18,20^. These conserved relationship between INM, mitosis position and delamination in two different epithelia presenting drastically different morphology and proliferation rates strongly suggest that it could be relevant in multiple configurations.

Apart from the acceleration of the cell cycle (particularly the G2/M phase), we found that a major factor promoting non-apical mitosis at the PS is relaxation of the actomyosin cytoskeleton. In silico simulations allowed to separately modify the tension in distinct cellular regions, which showed that apical actomyosin played little role in INM and was rather important for tissue stability. Indeed, the integrity and organization of F-actin is maintained in epiblast apical junctions throughout gastrulation, notably via ASPP2^45^, which prevents apical extrusion. In this context, it is important to note that apical mitoses fight against apical contractility^19^. Therefore, maintenance of normal INM at the PS would likely prevent apical constriction and the acquisition of the bottle shape morphology and subsequently lead to some apical extrusion.

Epiblast cells that ingress at the PS to become mesoderm adopt a bottle shape with a constricted apical surface^4,7^. Delamination is a progressive process that only involves a proportion of PS cells at a given time. The asynchronous shrinkage of apical junctions is regulated by a Crumbs2-regulated anisotropy of myosin II localization^7,8^. Similarly, cellular heterogeneity in the neural crest appears to boost epithelial destabilization and thereby favor delamination^20^.

In addition to the position of nuclei, polarized protrusive activity mediated by the interaction between α4 and 5 integrins in neural crest cells and the extracellular matrix promotes basal extrusion^20^. Likewise, nascent mesoderm displays numerous projections mostly oriented towards the visceral endoderm in the PS, then towards the front in the mesodermal wings. Those protrusions are required for directional migration, as demonstrated by the mesoderm migration defects identified in mouse embryos deficient for PTK7, Rac1, Nap1, or β-Pix^22,46–48^. Developing tissues go through phase transitions between an ordered state (solid-like) and a disorganized state (fluid-like), which occur at a critical point, the end of phase equilibrium^49^. During development, being in constant balance around this critical point may allow embryos to combine robustness and adaptability in order to go through morphogenetic events^49^. Both mitosis and EMT trigger changes in tissue visco-elastic properties that can participate in symmetry breaking, suggesting the applicability of rheology concepts to the epiblast of gastrulating mouse embryos^44^.

Through combining ex vivo and in silico experiments we found that gastrulation EMT coincides with relaxation of the cytoskeleton and change in cell cycle regulation at the primitive streak. These local changes disrupt the synchronization between apical nuclei movements and mitosis, leading to an elevated occurrence of non-apical mitoses and basal positioning of nuclei. This orchestration may ensure the proper progression of gastrulation while maintaining the integrity of the epithelium (Figure 7) and may be an evolutionarily conserved strategy common to other events involving EMT in pseudo-stratified epithelia^20^.

**Figure 7:**
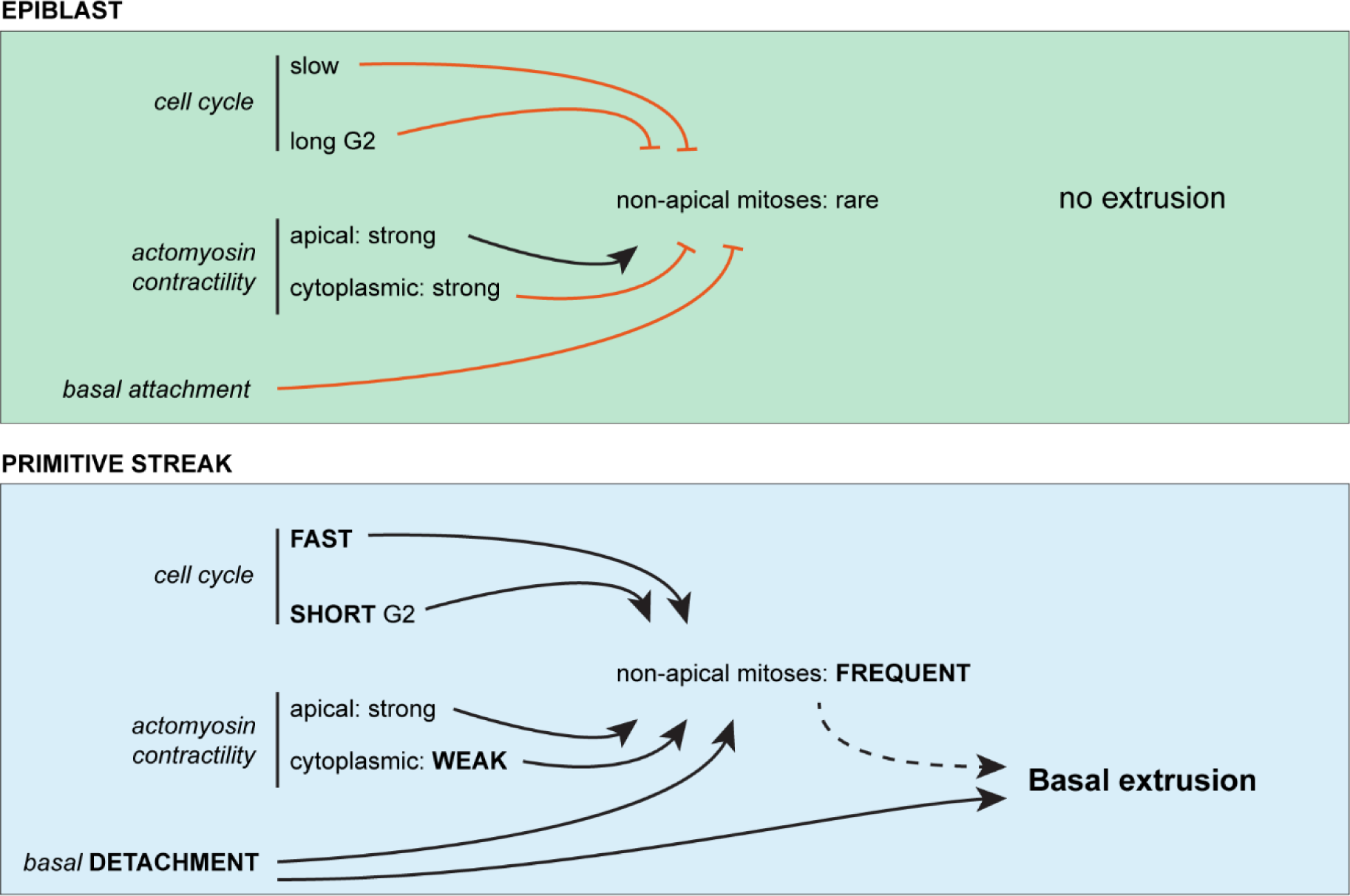
Occurrence of non-apical mitoses at the primitive streak induced by relaxation of actomyosin and acceleration of the cell cycle contributes to cell extrusion during mouse gastrulation. Diagram representing the state of the cell cycle, actomyosin contractility, and basal attachment in the region of the epiblast and the primitive streak during gastrulation EMT in the mouse embryo. At the primitive streak, faster cell cycle and shortened G2 duration, combined with weak cytoplasmic actomyosin contractility and basal detachment of cells, leads to a high frequency of non-apical mitosis that contribute to basal extrusion.

## MATERIAL AND METHODS

### Mouse breeding and genotyping

Mouse colonies were maintained in a certified animal facility in accordance with European guidelines. Experiments were approved by the local ethics committee (“Commission d’éthique et du bien-être animal”) under protocols 576N and 725N. Mouse lines were CD1 (Janvier Labs), mTmG ^50^, Sox2-Cre^41^, *RhoA*^38^, *ECT2*^39^, bred on a CD1 background; *CDK1*^40^mice were bred on a C57BL/6 background. Mouse genomic DNA was isolated from ear biopsies treated for 1 h at 95°C in NaOH to simultaneously genotype and identify animals.

For embryo genotyping, embryos were digested in Direct PCR mix (Viagen #102-T) with 1.5% Proteinase K (Quiagen #19133) for 2 hours at 56 C°, followed by 30 minutes at 90 C°.

The following sequences of primers were used for genotyping:

Sox2-cre: (F) TGCTGTTTCACTGGTTATGCTG, (R) TTGCCCCTGTTTCACTATCCAG. mTmG: (1) AAA GTC GCT CTG AGT TGT TAT, (2) GGA GCG GGA GAA ATG GAT ATG, (3) TCA ATG GGC GGG GGT CGT T.

RhoA fl: (1) AGC CAG CCT CTT GAC CGA TTT A, (2) TGT GGG ATA CCG TTT GAG CAT. RHOA KO: (1) AGG CAT GGA CCA CCA TGT CA, (2) ATG TGC TTC CCG TGT CTA GT. ECT FL: (1) GCA CTC CAA TTA TGA AGC (2) CAA TAT GTT GGG TAG AGA GAT GGC. ECT KO: (1) CAA TAT GTT GGG TAG AGA GAT GGC (2) TCC TCC GGG TGG ACC AGA G. CDK1 FL: (F) CCA GGG TGA CCT TGT CGT, (R) AGC CTG CCT CCA CTT CCA. CDK1 ko: (1) TTCTCCACGCTTGTCTCCAA (2) CAGCTTTAGGAGTGCAGGC.

### Antibodies

Antibodies for mouse embryo staining were: Goat anti-collagen IV (MERK #AB769, 1:500), rabbit anti-Phh3 (Sigma #SAB4504429, 1:500), rabbit anti-E-cadherin (Cell signaling #3195S, 1:500), rabbit anti-laminB1 (abcam #ab229025, 1:250), sheep anti N-cadherin (R&D #AF64216, 1/100), rabbit anti-vimentin (abcam #ab92547,1:200), goat anti-Brachyury (R&D #AF2085, 1/100), rabbit anti-caspase3 (R&D # AF835, 1:250), goat anti-snail (R&D #AF3639, 1:100). F-actin was visualized using rhodamine phalloidin (abcam #ab235138, 1:1000), and nuclei using DAPI (Sigma; 1:1000). Secondary antibodies were anti-goat Alexa Fluor 647 (Invitrogen #A21447, 1:500), anti-rabbit Alexa Fluor 488 (Invitrogen #A21206, 1:500), anti-sheep Alexa fluor 647 (Invitrogen #A11016, 1:500), anti-rabbit Alexa Fluor 647 (Jackson #711-605-152, 1:500) and anti-goat Alexa Fluor 488 (Invitrogen #A32814, 1:500).

### Embryo recovery, staging, and pharmacological treatment

Embryos were recovered at the appropriate time point after observation of a vaginal plug at day 0. E6.5 and E7.5 embryos were dissected in dissection medium using #5 forceps and tungsten needles under a transmitted light stereomicroscope. Dissection medium was composed of Dulbecco’s modified Eagle medium (DMEM) F-12 supplemented with 10mM HEPES and L-glutamin (Thermofisher, #11039047) with 10% Fetal Bovine Serum (Thermofisher, #10270106) and 1% Penicillin/Streptomycin (P/S). Bright-field pictures of the litter or single embryo were taken before any manipulation to ensure adequate staging. For pharmacological treatment, E6.5 embryos were allowed to recover in equilibrated culture medium (50% DMEM F12 (Thermofisher,# 21041025), 50% rat serum (Janvier), 1% P/S) in an incubator (37°C, 5% CO2) for 1 h. Embryos were then transferred to 15-wells ibidi (#81507) with 30 µl of equilibrated culture medium containing inhibitors or control reagent depending on what chemical the inhibitor was diluted in. Bright-field pictures were taken at t0 and after the incubation with inhibitors to assess embryo growth. For the post-treatment survival test, medium was replaced with fresh culture medium and embryos were cultured over night at 37C°.

Inhibitors were: PD16685 (Wee1 inhibitor, MECK #PZ0116, diluted in dH20), R03366 (CDK1 inhibitor, Sigma #217699 in DMSO), Y27632 (Rock inhibitor, Abcam #ab120129 in dH20), Prinomastat hydrochloride (Pan-MMPs inhibitor, Sigma #PZ198 in dH20), NSC405020 (MMP14 inhibitor, Tocris #4902/10 in ethanol).

### Immunofluorescence on mouse embryos

For immunofluorescence, embryos were fixed in PBS containing 4% paraformaldehyde (PFA) for 2h at 4°C. Embryos were cryopreserved in 30% sucrose, embedded in OCT and sectioned at 7µm. Staining was performed in PBS containing 0.5% Triton X-100, 0.1% BSA and 5% heat-inactivated horse serum. For whole mount, embryos were imaged in PBS 1X. Alternatively, to enhance optical clarity, E6.5 and E7.5 were treated in 35µl of Refraction Index Matching (RIM), as described in ^51^. For proximal-distal oriented whole-mount, embryos were embedded in warm 0.5% low melting agarose dissolved in PBS 1X, in a 15-well ibidi. Sections and whole-mount embryos were imaged on a Zeiss LSM 780 microscope equipped with Plan-Apochromat 25×/0.8, C Achroplan 32×/0.85 and LD C Apochromat 40×/1.1 objectives.

### Image analysis

Images were processed using Arivis Vision4D v2.12.3 (Arivis, Germany), image j Fiji and Icy software (http://icy.bioimageanalysis.org).

Mitotic index and non-apical mitotic index statistical analysis were performed as described in^12^. Mitosis was considered “non-apical” when happening at least 10 µm away from the apical pole and was not in the first pseudo-layer of nuclei lining the apical pole. Only cells still attached to the apical pole, and that did not cross the limit of basement membrane discontinuity, were considered in the measurements.

For transverse sections, anterior-posterior boundary was placed at mid-distance between the anterior and posterior poles. The PS region was defined by the area where the basement membrane was discontinuous, and the posterior region quantification excluded counts from the PS region. A cell was counted as being part of the PS region if at least 50% of its cell body was within the area where the basement membrane was non-ambiguously degraded, and if the cell retained its attachment to the apical pole (cell contours were defined by F-actin detection using Phalloidin). Phospho-histone H3 labelling was used to count cell in mitosis, normalised by the number of nuclei (DAPI staining) in each region. Mitosis was considered “non-apical” when happening at least 10 µm away from the apical pole and/or not in the first pseudo-layer of nuclei lining the apical pole. Non-apical mitosis ratio was calculated by dividing the number of non-apical Phh3+ nuclei by the total number of Phh3+ nuclei in a subregion of the epiblast. The percentage of basement degradation was calculated using Collagen IV labelling, through dividing the length of the area where the membrane was degraded by the total length of the basement membrane around the epiblast and conversion to percentage. Nuclei size ratio was calculated using Lamin B1 and DAPI labelling, by diving nuclei short axis by their long axis.

For each population, normality was assessed using a Shapiro– Wilk test. According to the results of the precedent test, samples were compared using a non-parametric Mann–Whitney test or an unpaired t-test. For cross statistics, either anova or Kruskall-Wallis followed by Dunn post tests were performed. Ns: non-significant, *P-value ≤ 0.05, **P-value ≤ 0.01 and ***P-value ≤ 0.001. Graphs and statistics were obtained using Spyder environment (3.9) python code and prism 5.0.

### In-Silico modeling

The complete details of the model are described in ^20^. Parameters used to adjust the in silico tissue to the epiblast configuration are mentioned in the Results section.

## ACKNOWLEDGMENTS

We wish to thank the Université Libre de Bruxelles/Erasme animal facility. We gratefully acknowledge the Université Libre de Bruxelles light microscopy (LiMiF) core facility (M. Martens and J-M. Vanderwinden) for help with confocal imaging.

E.D.G. received a FRIA fellowship of the Fonds de la Recherche Scientifique (FNRS) and a grant from the “Fondation Alice and David Van Buuren” and “Fondation Jaumotte-Demoulin”. I.M. is a FNRS senior research associate. E.T. is a research director at the French National Research Center (CNRS).

---

**Supplementary 1:**
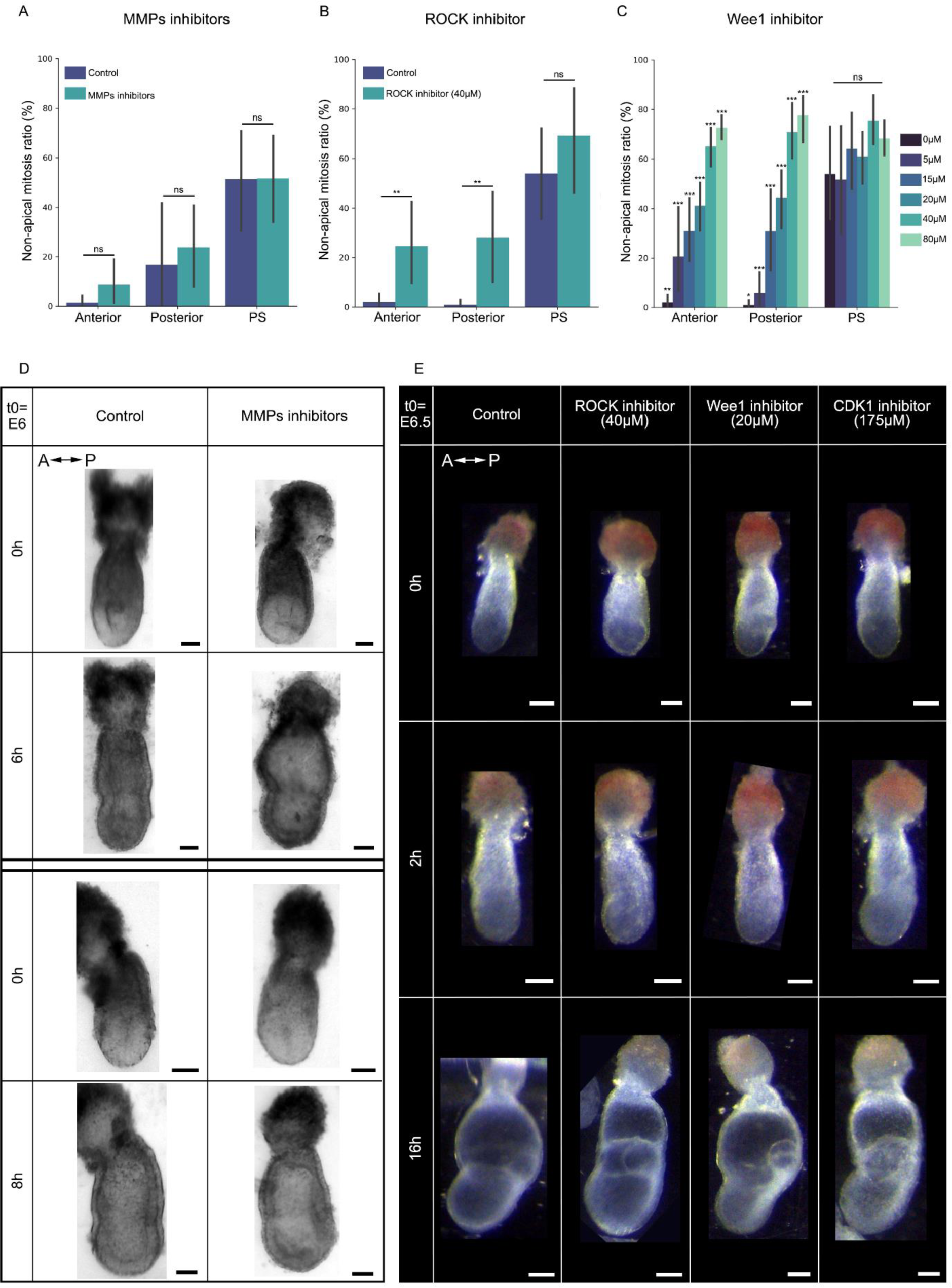
(A-C) Bar plots representing non-apical mitosis ratio in of embryos cultured in vehicle or for 6 hours with MMPs inhibitors (A); for 2 hours with 40µM of ROCK inhibitor (B); for 2 hours with 5µM, 10µM, 20µM, 40µM, 80µM of Wee1 inhibitor (C), in anterior, posterior and PS regions. Ns: nonsignificant, *: P-value≤0.05, **: P-value≤0.01 and ***: P-value≤0.001. Shapiro–Wilk test followed by Mann-Whitney or t-test. (A) Control: *n*= 14 slides from 8 embryos. MMPs inhibitors: *n*= 21 slides from 7 embryos. (B) Control: *n*= 26 slides from 12 embryos, ROCK inhibitor: *n*= 14 slides from 7 embryos. (C) Control: *n*= 27 slides from 13 embryos, 5µM Wee1 inhibitor: *n*= 12 slides from 5 embryos, 10µM Wee1 inhibitor: *n*= 16 slides from 10 embryos, 20µM Wee1 inhibitor: *n*= 21 slides from 9 embryos, 40µM Wee1 inhibitor: *n*= 9 slides from 4 embryos, 80µM Wee1 inhibitor: *n*= 12 slides from 6 embryos. (D) Brightfield images of control embryos and embryos cultured with MMPs inhibitors at t= 0h and t= 6 h (top) and t=0 h and t=8h (bottom). Scale bars: 50µm. 6 hours culture: control: *n=* 21; MMPs inhibitors: *n=* 29. 8 hours culture: control: *n=* 10, MMPs inhibitors: *n=* 9. (E) Brightfield images of E6.5 embryos vehicle, 40µM ROCK inhibitor, 20µM Wee1 inhibitor, and 175µM CDK1 inhibitor at t=0h, t=2h and t=16h. Embryos were cultured for two hours with the designated inhibitors then placed in fresh culture medium and cultured over night for a total of 16 hours post dissection. Scale bars: 50µm. Control: *n=* 10, 40µM ROCK inhibitor: *n=* 12, 20µM Wee1 inhibitor: *n=* 11, 175µM CDK1 inhibitor: *n=* 10. A-P: Anterior – Posterior.

**Supplementary 2:**
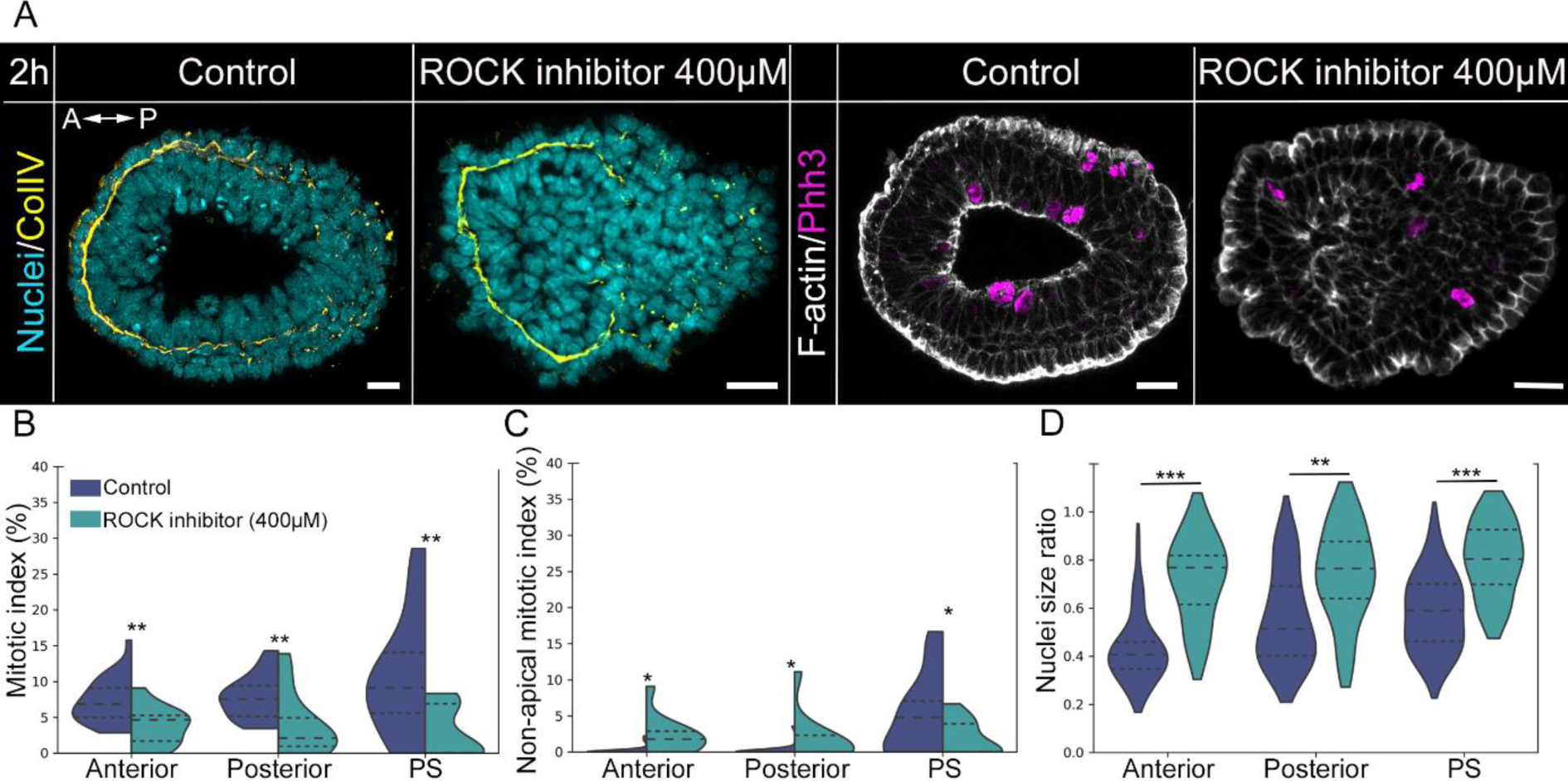
(A) Z-projection of transverse sections from E6.5 embryos cultured for 2 hours with control vehicle or ROCK inhibitor (Y27632) at 400µM, stained for nuclei (DAPI, cyan), basement membrane (collagen IV, yellow), mitosis (phh3, magenta) and F-actin (phalloidin, grey). Scale bars: 25µm. (C-C) Split violin plots representing the mitotic index (B) and non-apical mitotic index (F), in anterior, posterior, and PS regions in embryos cultured with control vehicle or 400µM ROCK inhibitor. Inner doted lines indicate third quartile, median, and first quartile. Ns: nonsignificant, *: P-value≤0.05, **: P-value≤0.01. Normality was assessed using a Shapiro–Wilk test followed by Mann-Whitney or t-test. Control: *n*= 26 slides from 12 embryos, anterior: 1048 nuclei, posterior: 663 nuclei, PS: 348 nuclei. ROCK inhibitor: *n*= 11 slides from 4 embryos, anterior: 620 nuclei, posterior: 439 nuclei, PS: 298 nuclei. (G) Violin plots representing nuclei size ratio in anterior, posterior, and PS regions, in embryos cultured with control vehicle or 400µM ROCK inhibitor. Inner doted lines indicate third quartile, median, and first quartile. Ns: non-significant, **: P-value≤0.01, ***: P-value≤0.001. Normality was assessed using a Shapiro–Wilk test followed by Mann-Whitney or t-test. Control: *n*= 11 embryos, anterior: 37 nuclei, posterior: 37 nuclei, PS: 37 nuclei. 400µM ROCK inhibitor: *n*= 4 embryos, anterior: 15 nuclei, posterior: 15 nuclei, PS: 15 nuclei.

**Supplementary 3:**
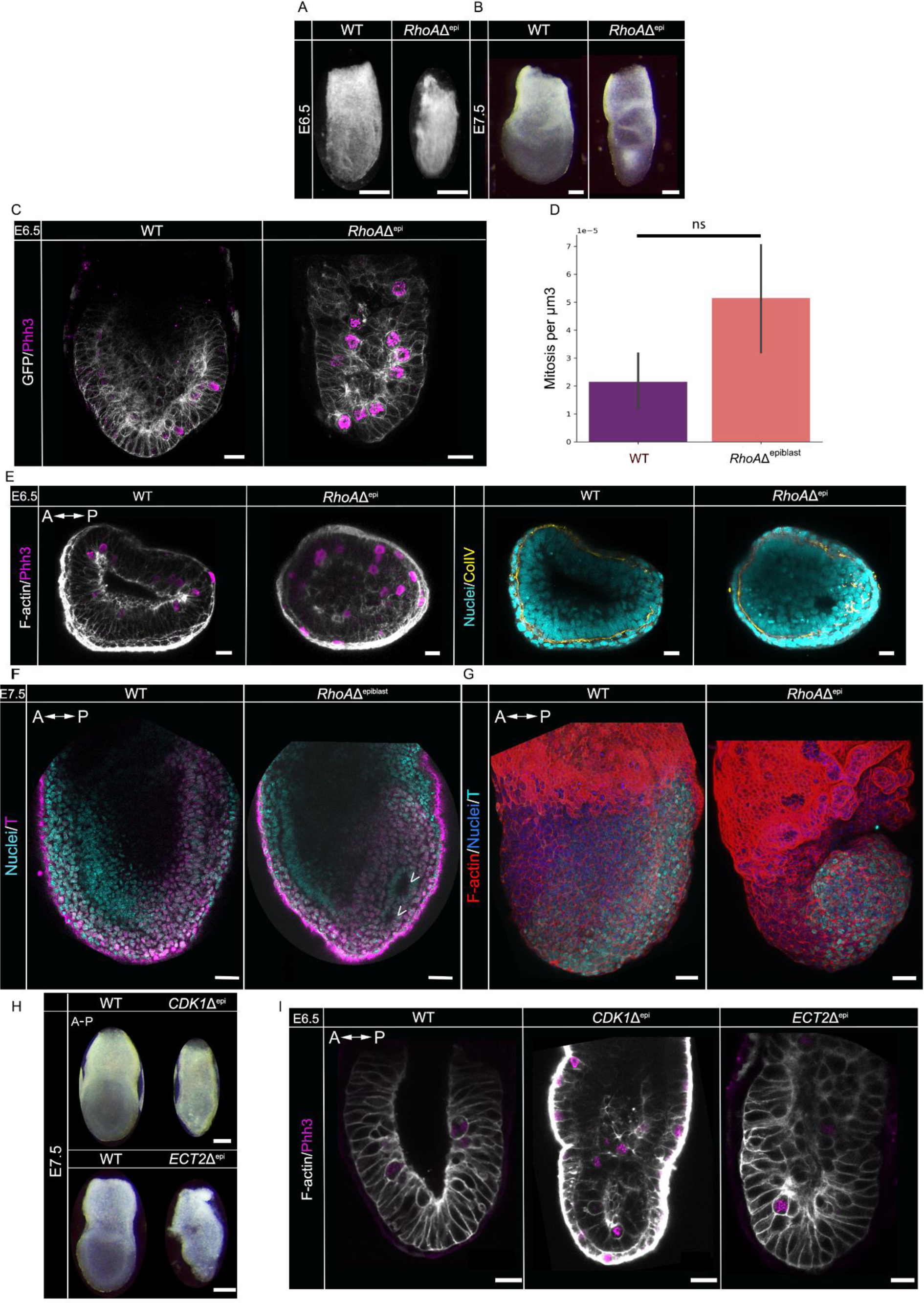
(A-B) Brightfield images of E6.5 (A) and E7.5 (B) wild-type (WT) and RhoAΔ^epiblast^ embryos. Scale bars: 100µm. (C) Z-projections in sagittal view from whole-mount E6.5 WT and RhoAΔ^epiblast^ embryos expressing membrane GFP in the epiblast (grey), stained for mitosis (Phh3, magenta). Scale bars: 25 µm. (D) Barplot representing the mean number of mitoses per µm3 in WT versus RhoAΔ^epiblast^ embryos. (E) Projections from selected z-slices of E6.5 whole-mount WT and RhoAΔ^epiblast^ embryos embedded in low melting agarose in a proximal-distal orientation. Embryos are stained for F-actin (phalloidin, grey), mitosis (Phh3, magenta), nuclei (DAPI, cyan) and basement membrane (collagen IV, yellow). Scale bars: 25µm. (F) Projections of selected Z slices from E7.5 whole-mount WT and RhoAΔ^epiblast^ embryos stained for nuclei (DAPI, blue) and T (magenta). Arrow heads indicates ectopic lumens. Scale bars: 50µm. (G) 3D projections of E7.5 WT and RhoAΔ^epiblast^ whole-mount embryos stained for F-actin (phalloidin, red), nuclei (DAPI, blue) and T (cyan). Scale bars: 50µm. (H) Brightfield images of E7.5 WT, CDK1Δ^epiblast^ and ECT2Δ^epiblast^ embryos. Scale bars: 100µm. CDK1Δ^epiblast^: *n*= 5, ECT2Δ^epiblast^: *n*= 6. (I) Z-projections from E6.5 WT, CDK1Δ^epiblast^ and ECT2Δ^epiblast^ embryos stained for F-actin (phalloidin, grey) and mitosis (phh3, magenta). Scale bars: 25µm. CDK1Δ^epiblast^: *n*= 4, ECT2Δ^epiblast^: *n*= 2.

